# Loop extrusion provides mechanical robustness to chromatin

**DOI:** 10.1101/2025.09.16.676638

**Authors:** Hossein Salari, Daniel Jost

## Abstract

Chromosomes are complex biopolymers folded into dynamic loops via a loop extrusion process and may experience various mechanical forces *in vivo*. We develop a force-dependent model of chromatin loop extrusion and investigate its mechanical consequences on chromosome organization using simulations and analytical theory. We show that loop extrusion alters the force–extension behavior of DNA in a non-monotonic manner: extrusion stiffens the chain at low forces but softens it at intermediate and high forces. Our model predicts hysteresis in pulling–recoiling cycles and out-of-equilibrium responses, consistent with recent single-chromosome stretching experiments. We further find that loop extrusion provides mechanical robustness to chromatin by promoting compaction while enabling rapid structural recovery after stress. These results establish loop extrusion as a key regulator of chromatin mechanics.

## Introduction

The three-dimensional (3D) organization of the genome plays a fundamental role in regulating essential cellular processes, including gene transcription [1], DNA replication [2], and repair [3, 4]. However, chromatin—the nucleoprotein polymer forming chromosomes—is not a static scaffold: it is dynamically remodeled and is frequently subjected to mechanical forces generated by various processes such as pulling centromeres by microtubule spindles during mitosis [5], cytoskeleton-mediated nuclear deformations that affect chromatin in the nuclear periphery [6], or the actions of different molecular motors on chromatin ssuch as DNA and RNA polymerases [7, 8]. These forces can impose significant tensile stresses on the genome [9]. How chromatin preserves its structural and functional integrity under such mechanical challenges remains a central question in genome biology.

A key mechanism driving 3D genome organization is DNA loop extrusion, an ATP-dependent process mediated by structural maintenance of chromosomes (SMC) complexes such as cohesin and condensin [10] that organizes chromatin into topologically associating domains (TADs) and regulatory loops [11, 12]. As the activity of such motors depends on the local forces acting on DNA [13, 14], loop extrusion may provide an adequate mechanism to adapt chromatin organization to mechanical stresses. For example, recent studies suggest that condensin I enhances the stiffness and stability of mitotic chromosomes [15], indicating a possible role for loop extrusion in resisting force-induced disruption. However, the interplay between chromatin mechanics and loop extrusion activity remains largely unexplored.

In this work, we present a polymer model of chromatin that incorporates force-dependent loop extrusion activity. Inspired by recent experiments *in vivo* and *in vitro* that examined the mechanical response of chromosomes under applied tension [16, 17], we systematically characterize the force-extension relation of loop extruded chromatin using simulations and polymer theory. This framework allows us to investigate how loop extrusion modulates the stiffness of the chromatin and contributes to its mechanical resilience.

### Force-dependent loop-extrusion polymer model

We developed a 3D polymer model of chromatin under tension (Supplementary Method). Following our previous work [18, 19], chromatin is modeled as a semi-flexible (Kuhn length *l*_*k*_ = 100 nm), self-avoiding chain. Each monomer, 20 nm in size, corresponds to 1 kb of genomic DNA (Fig. 1a) and evolves over time via standard kinetic Monte Carlo (MC) simulations, with one MC step equals ∼ 1 ms in real time.

**FIG. 1.**
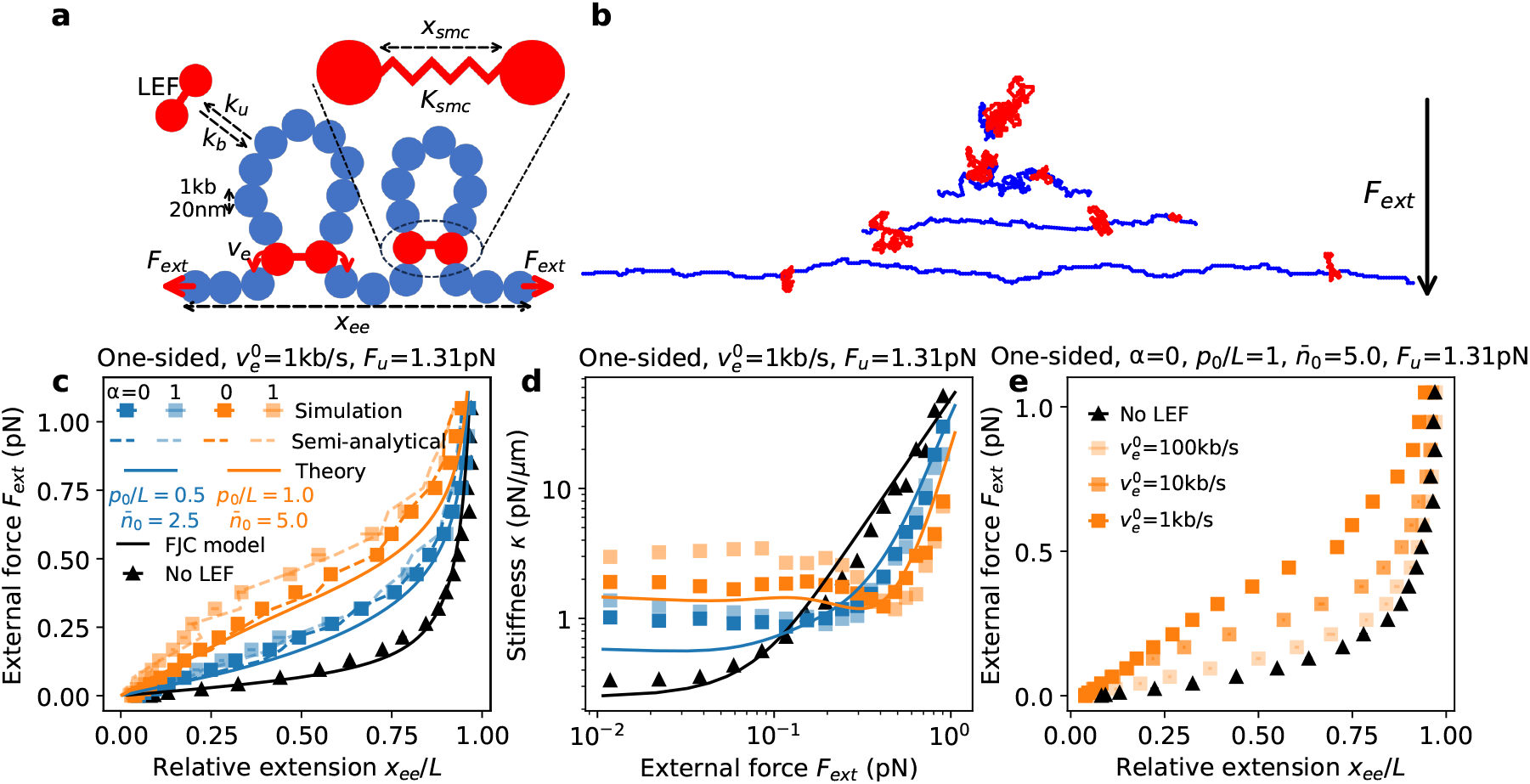
Anomalous force-extension curve induced by loop extrusion activity. **a)** Schematic representation of the polymer model under tension with SMC-mediated loop extrusion. **b)** Representative snapshots of the polymer under increasing external forces. Extruded segments are shown in red; non-extruded segments in blue. **c)** Force-extension curves for different loop extrusion activities. Symbols represent simulation results; curves denote theoretical predictions. **d)** Effective stiffness *κ* as a function of external force, extracted from the force-extension data shown in panel (c). Same color legend as in (c). **e)** Comparison of force-extension curves for various zero-force extrusion speeds 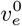 in the high activity regime and in the one-sided and *α* = 0 case. Symbols represent simulation results.

To model loop extrusion, we adopt a standard coarse-grained translocation scheme [18, 20]. Loop-extruding factors (LEFs) bind and unbind to chromatin at rates *k*_*b*_ and *k*_*u*_, respectively (Fig. 1a). At steady state, the probability of LEF binding is given by 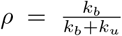, and the mean number of bound LEFs is 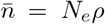, where *N*_*e*_ is the total number of LEFs. Upon binding, one LEF anchors its two “legs” to adjacent monomers and extrudes a loop by stochastically translocating the leg positions along the chain away from each other at a characteristic velocity *v*_*e*_. This movement can be one-sided or two-sided, depending on the extrusion mechanism. At steady state, the average residence time of a LEF on DNA is 1*/k*_*u*_, and its processivity, defined as the average extruded loop size in the absence of any barrier, is given by *p* = *v*_*e*_*/k*_*u*_.

To maintain physical connectivity between the LEF legs, we introduce a harmonic spring between them, of stiffness *K*_smc_. Legs of different LEFs may pass through one another depending on a crossing probability *α*, ranging from 0 (no possible crossing) to 1 (phantom legs).

Translocation of a leg halts when it encounters a boundary element.

To simulate mechanical stretching, equal and opposite external forces of strength *F*_*ext*_ are applied at the two opposite ends of the chain.

Within this framework, the local force exerted on a LEF at a given time, *F*_smc_, is: *F*_smc_ = *K*_smc_(*x*_smc_ − *x*_0_), where *x*_smc_ is the current spatial separation between the two legs, and *x*_0_ = 20 nm is the rest length of the spring. We choose *K*_smc_ = 0.031 pN/nm consistent with recent *in vitro* experiments estimating that a displacement of ∼32 nm between cohesin’s head and hinge domains (the two possible legs) generates a restoring force of ∼1pN [21, 22].

Many *in vitro* experiments on naked DNA have shown that increasing the mechanical forces acting on the extruded DNA template slows down the loop extrusion activity[13, 14]. To incorporate this effect, we assume that the extrusion velocity decreases exponentially with the force felt by the LEF:

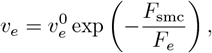

where 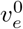 is the extrusion velocity in the absence of force, and *F*_*e*_ is the characteristic force-scale determining the sensitivity of LEFs to mechanical load. By fitting this model to available experimental data on the force-dependency of the velocity of condensin [14] (Supplementary Figure S1), we found 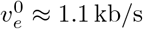 and *F*_*e*_ ≈ 0.2 pN. Notably, the value of *F*_*e*_ aligns well with the reported stall force of condensin [14].

We also extend this reasoning to LEF unbinding kinetics. As DNA tension may destabilize protein-DNA interactions, we assume that the unbinding rate increases exponentially with force:

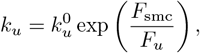

where 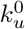 is the tension-free unbinding rate, and *F*_*u*_ is the force-scale governing unbinding sensitivity. Assuming that LEF dissociation is less sensitive to force than extrusion (*F*_*u*_ *> F*_*e*_), we explore a range of *F*_*u*_ values: 0.65, 1.31, and 13.1 pN.

With this model, we investigate the behavior of a 500 kb-long chromatin segment with fixed boundary elements at each end, at steady-state. The applied external force is systematically varied quasi-statically from 0 to 1 pN. Two loop extrusion regimes are considered: a “low”-activity regime with zero-force processivity 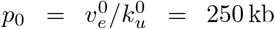 and mean LEF number 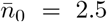, and a “high”-activity regime with *p*_0_ = 500 kb and 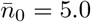. For both regimes, we simulate both one-sided (only one leg, randomly chosen, can translocate at velocity *v*_*e*_) and two-sided (both legs can translocate, each at velocity *v*_*e*_*/*2) loop extrusion scenarios for *α* = 0 and *α* = 1. Additionally, we vary the zero-force extrusion velocity 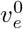 across a wide range—[1.0, 10, 100] kb/s—to systematically explore how loop extrusion speed may influence chromatin mechanics.

### Loop extrusion alters chromatin mechanics by tuning stored length

Overall, simulations reveal that the decrease of LEF speed and dwell time under tension leads to a reduction in the average loop sizes and numbers as well as in the fraction of DNA engaged inside loops (Fig. 1b and Supplementary Figure S2).

To assess the mechanical impact of loop extrusion, we derive the force–extension (FE) curves of the polymer for each investigated set of parameters (Fig. 1c and Supplementary Figure S3, colored symbols). In all cases, the FE curve differs qualitatively from the situation without LEFs (black symbols), exhibiting a characteristic “tilde-like” shape. In particular, for a given applied force, the extruded polymer exhibits a lower extension because of the compaction mediated by loop formation. This compaction is more pronounced in the “high”-activity regime and is slightly stronger for two-sided extrusion than for one-sided. Moreover, the effect is amplified when LEF legs are allowed to cross one another (*α* = 1 vs *α* = 0). Finally, we find that increasing the unbinding force scale *F*_*u*_ enhances this behavior (Supplementary Figure S4).

We show that the FE behavior of a chain without loop extrusion is accurately captured by the Freely-Jointed Chain (FJC) model with *l*_*k*_ = 100nm (Fig. 1c). Building on this agreement, we extend the FJC framework to incorporate the effects of loop extrusion. The extension (projection of end-to-end distance along the force direction) *x*_ee_ of a FJC under tension is given by [23]:

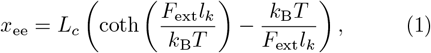

where *L*_*c*_ is the contour length of the polymer, *k*_B_ is the Boltzmann constant, and *T* is the absolute temperature. In the presence of active loop extrusion, a fraction of the DNA is dynamically sequestered within loops, reducing the effective contour length 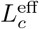 available for extension. From simulations, we can estimate directly 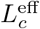 and then predict the extension via the FJC model (dashed curves in Fig. 1c). This semi-analytical approach demonstrates that the primary effect of loop extrusion on the FE curve can be interpreted as a reduction in the effective contour length.

To obtain a theoretical prediction for 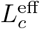, we consider a chromatin domain of length *L* with 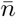 LEFs, either two-sided or one-sided. For two-sided LEFs, we assume that the domain is partitioned into 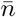 equal subdomains of size 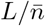, with each LEF acting independently within its subdomain. For one-sided LEFs, the effective number of extruding legs is halved, so the domain is divided into 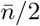 subdomains of size 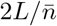. In this framework, the average loop size extruded by a single LEF can be approximated as (Supplementary Method):

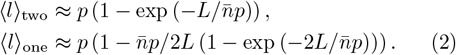

The total fraction of DNA engaged within extruded loops can then be estimated by multiplying the number of LEFs by the average loop size.

Since both *p* and 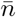 are force dependent, we assume the applied external force *F*_ext_ is uniformly transmitted along the chain, and each LEF experiences this force. Accordingly, we express the force-dependent processivity and LEF occupancy as:

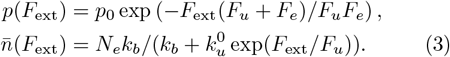

Our analytical predictions qualitatively recapitulate the simulation results in both the “low” and “high” loop-extrusion activity regimes (Fig. 1c).

### Non-monotonic force-dependent chromatin stiffness

We further evaluate the stiffness of the polymer under loop extrusion, defined as *κ* = *∂F*_ext_*/∂x*_ee_ (Fig. 1d). For an ideal FJC without loop extrusion, stiffness exhibits two regimes: at low forces, *κ* is approximately constant, while at high forces, 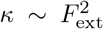. When loop extrusion is active, we observe an enhancement of stiffness in the low-force regime, which becomes more pronounced with increasing extrusion activity. This behavior reflects the tendency of LEFs to resist chain extension by compacting the polymer into loops. In contrast, for larger applied forces, above *F*_*e*_, the stiffness becomes lower than that of the ideal chain. Interestingly, the force–stiffness relationship becomes non-monotonic: *κ* reaches a local minimum at an intermediate force regime before rising again at high forces. This behavior is especially pronounced for systems with “high” loop extrusion activity. Indeed, when loops start to be disrupted near the stall force, the sudden release of the monomers stored within the loops leads to a sharp increase in the effective contour length and thus a transient softening of the polymer [24]. Finally, at very high forces, all stiffness curves converge to the ideal chain behavior, indicating that LEFs become effectively inactive under strong tension.

### Impact of loop extrusion speed on chromatin mechanics

Next, we investigate how the zero-force loop extrusion speed 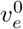 influences the effective stiffness of the polymer (Fig. 1e) at fixed zero-force processivity *p*_0_ and average number of LEFs 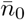. We find that the impact of loop extrusion on the FE diminishes as the extrusion speed 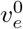 increases. Despite maintaining constant initial processivity and binding probability, both the average number and size of extruded loops decrease with increasing extrusion speed (Supplementary Figure S5). These findings indicate that the characteristic time scale of loop extrusion activity, *t*_*e*_ ∼ 1kbp*/v*_*e*_, becomes shorter than the polymer relaxation time, which is approximately *t*_*p*_ ≈ 0.5*s* in our simulations (estimated from the relaxation dynamics of the end-to-end distance of a 100-kbp DNA, Supplementary Figure S6). As a result, the legs of the LEF may not have sufficient time to reach their equilibrium configuration, leading to transiently elevated internal forces. These non-equilibrium forces can increase the unbinding rate and reduce the effective extrusion rate, thereby lowering the functional processivity. Consequently, fast-extruding LEFs may not significantly deform the chromatin before detaching, thereby limiting their contribution to the overall mechanical response of the polymer.

### Loop extrusion enhances chromatin stability under tension

To explore how loop extrusion stabilizes chromatin under mechanical tension, we compute the contact probability within a chromatin domain subjected to varying external forces (Fig. 2a and Supplementary Figure S7a). In the absence of loop extrusion, even weak forces (as low as ∼0.1 pN) lead to a dramatic loss of contacts, indicating significant conformational changes. In contrast, when loop extrusion is active, the polymer structure remains largely unaffected up to forces of approximately 0.3 pN.

**FIG. 2.**
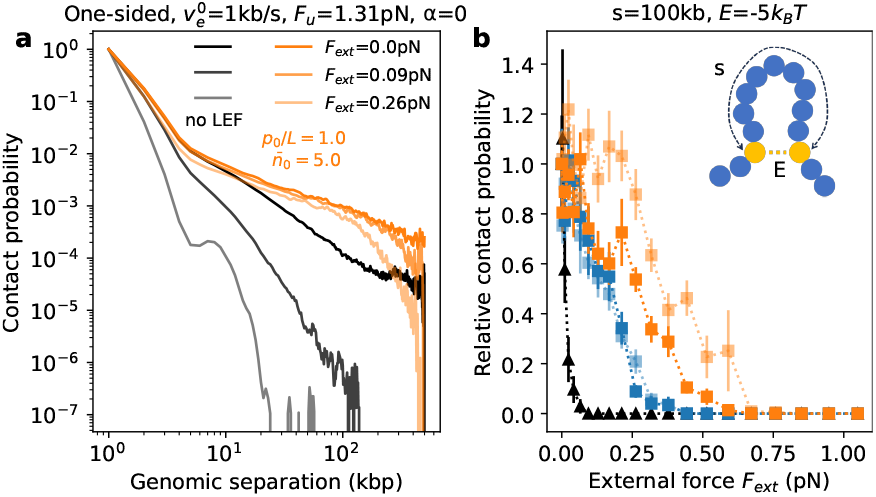
Polymer structure stabilization by loop extrusion. **a)** Contact probability as a function of genomic separation for different loop extrusion activities and external forces. **b)** Relative contact probability between two sticky monomers separated by 100 kb, plotted as a function of external force. The relative contact probability is defined as the contact probability normalized by its value at zero force. The legend follows the same convention as in Fig. 1c.

To further assess structural stability, we introduce a pair of “sticky” monomers—representing regulatory elements such as an enhancer and a promoter—separated by 100 kb and interacting via a short-range attraction of −5 *k*_B_*T* (Fig. 2b and Supplementary Figure S7b). We evaluate how the contact probability between these sites is altered by increasing external force, both in the presence and absence of loop extrusion. Without loop extrusion, the contact probability between these monomers drops below 5% of its zero-force value at just 0.1 pN. With active loop extrusion, however, this contact remains largely preserved at the same force level. To quantify this stabilizing effect, we define the rupture force as the external force required to reduce the contact probability to 5% of its zero-force value. The rupture force is approximately ∼0.1 pN without loop extrusion, increases to ∼0.4 pN under “low” extrusion activity, and reaches about ∼0.7 pN in the “high”-activity regime. These results demonstrate that loop extrusion significantly enhances the mechanical resilience of chromatin, helping maintain structural integrity under external forces.

### Loop extrusion controls chromatin relaxation and recovery under stress

We then examined the dynamic response of the chromatin chain under mechanical load using stress-relaxation simulations. After equilibrating the system, we applied a sudden increase in external tension, maintained the applied force for 1000 seconds, and then abruptly released it to zero (Fig. 3a). By tracking the chain extension over time, we observed a classical viscoelastic response that is well described by a linear Kelvin–Voigt model (Fig. 3a insets). During the pulling phase, the end-to-end extension follows an exponential approach to its steady-state value:

**FIG. 3.**
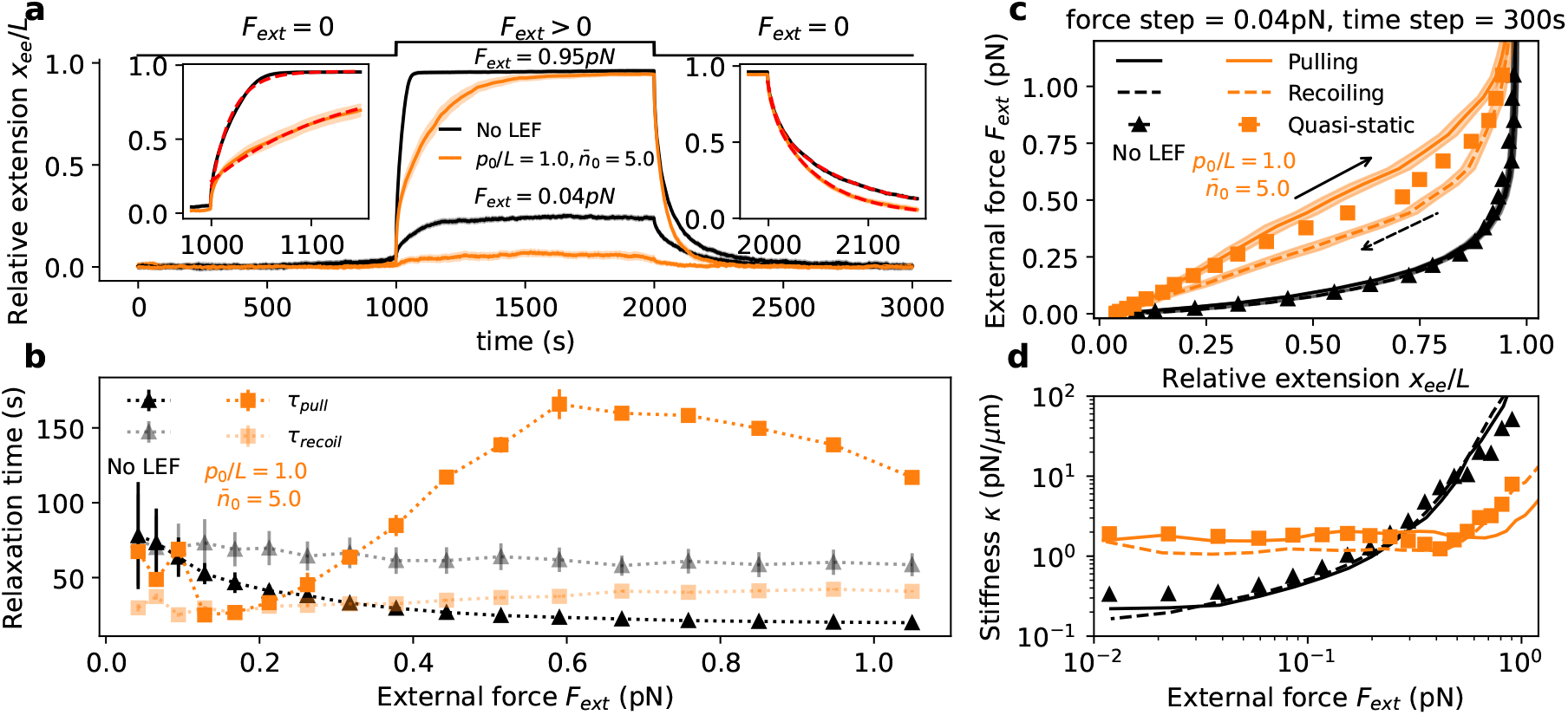
Dynamic response of a polymer with loop extrusion under external force. **a)** Relative extension as a function of time for different levels of loop extrusion activity. The external force is held at zero from 0 to 1000 s, then suddenly increased at *t*_1_ = 1000 s and maintained constant until *t*_2_ = 2000 s, after which it is abruptly set back to zero and held from 2000 s to 3000 s. Two representative force magnitudes are shown: low force (0.04 pN) and high force (0.95 pN). Insets highlight the extension dynamics during the transition periods in the high-force case, with exponential fits shown as red dashed lines. b) Characteristic relaxation times during pulling and release phases, obtained from exponential fits, plotted as a function of applied force for varying loop extrusion activities. **c)** Force–extension curves for both pulling and recoiling phases under different loop extrusion activities, when the applied tension is changed by discrete steps of 0.04 pN every 300 seconds, highlighting the emergence of hysteresis. **d)** Effective stiffness as a function of external force, derived from the curves in panel (c).

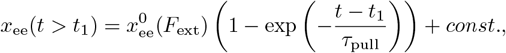

where *t*_1_ is the pulling start time, 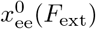 is the steady-state extension under applied force, and *τ*_pull_ is the characteristic relaxation time during force application. Upon force removal at time *t*_2_, the chain recoils exponentially:

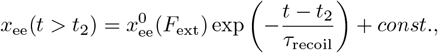

where *τ*_recoil_ denotes the relaxation time in the unloading phase. Both relaxation times are inversely related to the effective stiffness of the chain, i.e., *τ*_pull_, *τ*_recoil_ ∼ 1*/κ*.

In the absence of loop extrusion, we find that *τ*_recoil_ *> τ*_pull_, indicating that the chain responds more quickly to the application of force than to its release (Fig. 3b). Moreover, *τ*_pull_ decreases with increasing external tension, reflecting a force-induced stiffening of the chromatin chain and a nonlinear viscoelastic response. In contrast, *τ*_recoil_ remains largely independent of the applied force, suggesting that it is governed by the zero-force stiffness of the elastic chain and the intrinsic relaxation dynamics of the entire chain.

When loop extrusion is active, this behavior is reversed: we observe *τ*_pull_ *> τ*_recoil_ (Fig. 3b). Furthermore, *τ*_pull_ exhibits a non-monotonic dependence on force: increasing at low to intermediate forces, then decreasing at higher forces. This trend becomes more pronounced with higher loop extrusion activity (Supplementary Figure S8) and reflects the non-monotonic force-stiffness relationship observed in extruding chains. Notably, *τ*_recoil_ is reduced relative to the non-extruding case, reflecting a higher stiffness due to loop-induced compaction in the absence of external load.

At high forces (≳0.2 pN), *τ*_pull_ in the presence of loop extrusion exceeds that in the absence of loop extruders. This suggests a delayed deformation response due to structural constraints imposed by persistent loops. In contrast, the chain relaxes more rapidly when the force is removed, highlighting the system’s ability to efficiently return to compact configurations. These results suggest that loop extrusion protects chromatin integrity under load by resisting deformation and promoting rapid structural recovery, effectively preserving the chromatin conformation under mechanical stress.

### Hysteresis in extruded chromatin under tension

The out-of-equilibrium, dynamical response of extruded chromatin can also be observed by studying the FE behavior but in a non-quasi-static way, i.e., without allowing the system to reach a full steady-state while increasing or decreasing the applied tension (Fig. 3c and Supplementary Figure 9). A hysteresis cycle is observed, the system keeping the memory of the previous more compact or more elongated state when the external force is gradually augmented or reduced, respectively. Interestingly, the non-monotonicity of the effective stiffness *κ* during the pulling and recoiling phases is much less pronounced, and the polymer is stiffer when pulling up to high forces (∼ 0.7 pN) (Fig. 3d).

### Conclusions

We have introduced a force-dependent loop extrusion model and demonstrated its profound impact on chromatin mechanics. Our key finding is that loop extrusion significantly alters the stiffness of chromatin in a force-dependent and non-monotonic manner: extrusion stiffens the chromatin at low forces but softens it at intermediate and high forces. This behavior is characteristic of the force-extension curve of polymers with internal stored-length [24], such as RNA molecules whose internal secondary structures may unzip at high forces [25]. Our analytical model suggests that this effect is dominated by the force-dependent changes in the effective contour length of the chain. Our predictions are qualitatively in agreement with recent *in vitro* force-extension experiments on human mitotic chromosomes [15, 17], which are known to be structured by condensin-mediated loop extrusion [26]. Indeed, they observed hysteresis force-extension behavior and a force-dependent stiffness consistent with our out-of-equilibrium pulling/recoiling simulations, in particular with a softer stiffness at high forces.

We show that loop extrusion not only compacts chromatin structurally but also enhances its mechanical robustness—resisting deformation under sudden stress and promoting rapid structural recovery. This dual role may be critical for maintaining chromosome architecture and safeguarding essential cellular processes, such as transcription and replication, against mechanical perturbations. Our findings suggest new experimental directions to explore how chromatin responds to mechanical stress *in vivo* [16], particularly in cells lacking or overexpressing key loop-extruding factors.

## Acknowledgments

We are grateful to Chunlong Chen, John Marko and members of the Jost group for fruitful discussions. We acknowledge Agence Nationale de la Recherche (Grants No. ANR-21-CE45-0011, ANR-22-CE12-0035) for funding. We thank Centre Blaise Pascal de simulation et modélisation numérique of the ENS de Lyon for computing resources.

## Code Availability

The Fortran code implemented in this study is openly accessible via GitHub: https://github.com/physical-biology-of-chromatin/ForceExtensionLoopExtrusion.

## References

[1] J. Zuin, G. Roth, Y. Zhan, J. Cramard, J. Redolfi, E. Piskadlo, P. Mach, M. Kryzhanovska, G. Tihanyi, H. Kohler, et al., Nature 604, 571 (2022).

[2] D. J. Emerson, P. A. Zhao, A. L. Cook, R. J. Barnett, K. N. Klein, D. Saulebekova, C. Ge, L. Zhou, Z. Simandi, M. K. Minsk, et al., Nature 606, 812 (2022).

[3] A. Piazza, H. Bordelet, A. Dumont, A. Thierry, J. Savocco, F. Girard, and R. Koszul, Nature cell biology 23, 1176 (2021).

[4] C. Arnould, V. Rocher, F. Saur, A. S. Bader, F. Muzzopappa, S. Collins, E. Lesage, B. Le Bozec, N. Puget, T. Clouaire, et al., Nature 623, 183 (2023).

[5] B. Akiyoshi, K. K. Sarangapani, A. F. Powers, C. R. Nelson, S. L. Reichow, H. Arellano-Santoyo, T. Gonen, J. A. Ranish, C. L. Asbury, and S. Biggins, Nature 468, 576 (2010).

[6] Y. Kalukula, A. D. Stephens, J. Lammerding, and S. Gabriele, Nature reviews Molecular cell biology 23, 583 (2022).

[7] B. Maier, D. Bensimon, and V. Croquette, Proceedings of the National Academy of Sciences 97, 12002 (2000).

[8] H. Yin, M. D. Wang, K. Svoboda, R. Landick, S. M. Block, and J. Gelles, Science 270, 1653 (1995).

[9] D. Saintillan, M. J. Shelley, and A. Zidovska, Proceedings of the National Academy of Sciences 115, 11442 (2018).

[10] C. Dekker, C. H. Haering, J.-M. Peters, and B. D. Rowland, Science 382, 646 (2023).

[11] S. S. Rao, S.-C. Huang, B. G. St Hilaire, J. M. Engreitz, E. M. Perez, K.-R. Kieffer-Kwon, A. L. Sanborn, S. E. Johnstone, G. D. Bascom, I. D. Bochkov, et al., Cell 171, 305 (2017).

[12] M. Gabriele, H. B. Brandão, S. Grosse-Holz, A. Jha, G. M. Dailey, C. Cattoglio, T.-H. S. Hsieh, L. Mirny, C. Zechner, and A. S. Hansen, Science 376, 496 (2022).

[13] I. F. Davidson, B. Bauer, D. Goetz, W. Tang, G. Wutz, and J.-M. Peters, Science 366, 1338 (2019).

[14] M. Ganji, I. A. Shaltiel, S. Bisht, E. Kim, A. Kalichava, C. H. Haering, and C. Dekker, Science 360, 102 (2018).

[15] C. F. Nielsen, H. Witt, A. Ridolfi, B. Kempers, E. M. Chameau, S. van der Smagt, M. Barisic, E. J. Peterman, G. J. Wuite, and I. D. Hickson, bioRxiv, 2025 (2025).

[16] V. I. Keizer, S. Grosse-Holz, M. Woringer, L. Zambon, K. Aizel, M. Bongaerts, F. Delille, L. Kolar-Znika, V. F. Scolari, S. Hoffmann, et al., Science 377, 489 (2022).

[17] A. E. Meijering, K. Sarlos, C. F. Nielsen, H. Witt, J. Harju, E. Kerklingh, G. H. Haasnoot, A. H. Bizard, I. Heller, C. P. Broedersz, et al., Nature 605, 545 (2022).

[18] H. Salari, M. Di Stefano, and D. Jost, Genome research 32, 28 (2022).

[19] H. Salari, G. Fourel, and D. Jost, Nature Communications 15, 5393 (2024).

[20] G. Fudenberg, M. Imakaev, C. Lu, A. Goloborodko, N. Abdennur, and L. Mirny, Cell Rep. 15, 2038 (2016).

[21] G. Pobegalov, L.-Y. Chu, J.-M. Peters, and M. I. Molodtsov, Nature Communications 14, 3946 (2023).

[22] A. Bonato, J.-W. Jang, D.-G. Kim, K.-W. Moon, D. Michieletto, and J.-K. Ryu, Nucleic Acids Research 53, gkaf725 (2025).

[23] S. B. Smith, L. Finzi, and C. Bustamante, Science 258, 1122 (1992).

[24] S. Cocco, J. Marko, R. Monasson, A. Sarkar, and J. Yan, The European Physical Journal E 10, 249 (2003).

[25] P. Rissone, C. V. Bizarro, and F. Ritort, Proceedings of the National Academy of Sciences 119, e2025575119 (2022).

[26] K. Samejima, J. H. Gibcus, S. Abraham, F. Cisneros-Soberanis, I. Samejima, A. J. Beckett, N. Pučeková, M. A. Abad, C. Spanos, B. Medina-Pritchard, et al., Science 388, eadq1709 (2025).

