## Supplementary Information for "Loop extrusion provides mechanical robustness to chromatin"

### Supplementary information: Loop extrusion provides mechanical robustness to chromatin

Hossein Salari[1, 2, *] and Daniel Jost[1, †]
[1]*Laboratoire de Biologie et Modélisation de la Cellule, École Normale Supérieure de Lyon, CNRS, UMR5239, Inserm U1293, Université Claude Bernard Lyon 1, 46 Allée d'Italie, 69007 Lyon, France*
[2]*Institut Curie, PSL Research University, CNRS UMR3244, Dynamics of Genetic Information, Sorbonne Université, 75005 Paris, France*
(Dated: September 16, 2025)

#### SUPPLEMENTARY METHOD

##### A. Polymer model

We model the chromatin as a polymer chain undergoing local movements on a face-centered cubic (FCC) lattice with periodic boundary conditions, using the Metropolis algorithm as in our previous work [1, 2]. The total Hamiltonian of a given configuration is given by:

$$H = \kappa_b \sum_i (1 - \cos\theta_i) + \frac{1}{2} K_{\mathrm{smc}} \sum_{i<j} f_{ij} (x_{ij} - x_0)^2 - F_{\mathrm{ext}} x_{\mathrm{ee}},$$

where the first term describes the bending energy of the chain, with $\kappa_b$ denoting the bending rigidity and $\theta_i$ the angle between successive bond vectors at monomer $i$. The second term accounts for the harmonic potential between the two legs of a LEF, where $x_{ij}$ is the distance between monomers $i$ and $j$, and $f_{ij} = 1$ if monomers $i$ and $j$ are connected by a LEF, and 0 otherwise. The third term models the work done by an external force $F_{\mathrm{ext}}$, where $x_{\mathrm{ee}}$ is the extension along the force direction.

##### B. Analytical solutions for average extruded loop size

In the simplest case—where a single LEF operates on an infinitely long DNA molecule without boundaries or interactions with other LEFs (i.e., $\alpha = 1$)—the distribution of loop sizes $l$ follows a simple exponential form [3]:

$$P(l) = \frac{1}{p} \exp\left(-\frac{l}{p}\right). \tag{1}$$

When loop extrusion occurs within a confined domain of length $L$, bounded by two barrier elements, the loop size distribution differs depending on whether extrusion is two-sided or one-sided.

**Two-sided LEFs.** For two-sided LEFs, the probability distribution of loop sizes is approximated by:

$$P_{\mathrm{two}}^{\alpha=1}(l) \approx \begin{cases} \frac{1}{p} \exp\left(-\frac{l}{p}\right), & \text{for } l < L, \\ \exp\left(-\frac{L}{p}\right), & \text{for } l = L. \end{cases} \tag{2}$$

This distribution displays a noticeable peak at $l = L$, corresponding to LEFs that reach the domain boundary before dissociating. For simplicity, we neglect finite-size boundary effects on loops smaller than $L$, even though one leg of the LEF may transiently encounter the boundary during extrusion.

Under these assumptions, the average loop size is given by:

$$\langle l \rangle_{\mathrm{two}}^{\alpha=1} \approx p\left(1 - \exp\left(-\frac{L}{p}\right)\right). \tag{3}$$

This expression exhibits two limiting behaviors: For large domains ($L \gg p$), $\langle l \rangle \to p$, recovering the result for an unconstrained system. For small domains ($L \ll p$), extrusion is typically stopped by the boundaries, giving $\langle l \rangle \to L$.

**One-sided LEFs.** For one-sided LEFs operating within a domain of size $L$, the loop size distribution becomes:

$$P_{\mathrm{one}}^{\alpha=1}(l) = \frac{L + p - l}{pL} \exp\left(-\frac{l}{p}\right). \tag{4}$$

Unlike the two-sided case, the one-sided distribution does not exhibit a sharp peak at $l = L$ due to the asymmetric nature of extrusion. The corresponding average loop size is:

$$\langle l \rangle_{\mathrm{one}}^{\alpha=1} = p\left(1 - \frac{p}{L}\left(1 - \exp\left(-\frac{L}{p}\right)\right)\right). \tag{5}$$

In the two limiting cases: For $L \gg p$, $\langle l \rangle \to p$, consistent with unconstrained behavior. For $L \ll p$, only one leg reaches the boundary before unbinding, and $\langle l \rangle \approx L/2$.

Across all domain sizes, one-sided LEFs generate shorter average loops than two-sided LEFs due to their directional asymmetry.

Next, we consider the non-phantom case, where LEFs cannot cross each other.

**Two-sided LEFs.** For $\bar{n}$ non-phantom two-sided LEFs in a domain of size $L$, we approximate the system by dividing the domain into $\bar{n}$ equal segments, with one LEF per segment. The average loop size is:

$$\langle l \rangle_{\text{two}}^{\alpha=0} \approx p \left(1 - \exp\left(-\frac{L}{\bar{n}p}\right)\right). \tag{6}$$

**One-sided LEFs.** For one-sided non-phantom LEFs, each LEF effectively occupies half of a subdomain due to asymmetric extrusion. Thus, the system is modeled as $\bar{n}/2$ LEFs extruding within domains of size $2L/\bar{n}$, leading to the approximation:

$$\langle l \rangle_{\text{one}}^{\alpha=0} \approx p \left(1 - \frac{\bar{n}p}{2L}\left(1 - \exp\left(-\frac{2L}{\bar{n}p}\right)\right)\right). \tag{7}$$


*

†



[1] H. Salari, M. Di Stefano, and D. Jost, Genome research **32**, 28 (2022).

[2] H. Salari, G. Fourel, and D. Jost, Nature Communications **15**, 5393 (2024).

[3] C. A. Brackley, J. Johnson, D. Michieletto, A. N. Morozov, M. Nicodemi, P. R. Cook, and D. Marenduzzo, Physical review letters **119**, 138101 (2017).

**SUPPLEMENTARY FIGURES**

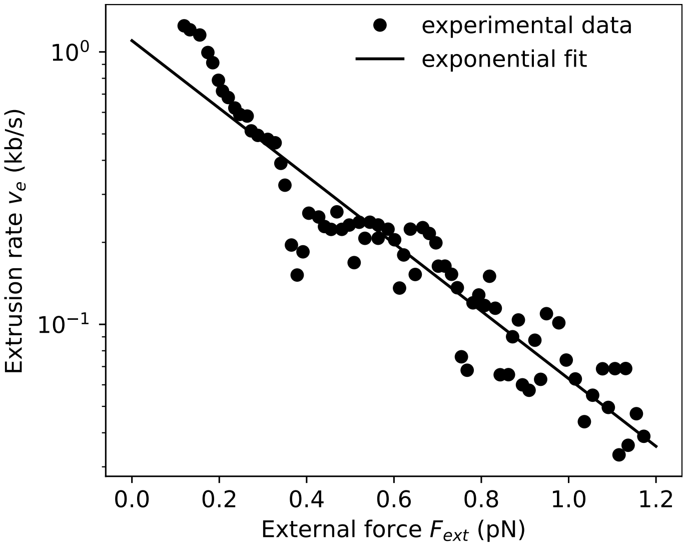


FIG. S1. Comparison between experimental measurements of loop extrusion rate as a function of external force (Ganji *et al.*) and the corresponding exponential fit.

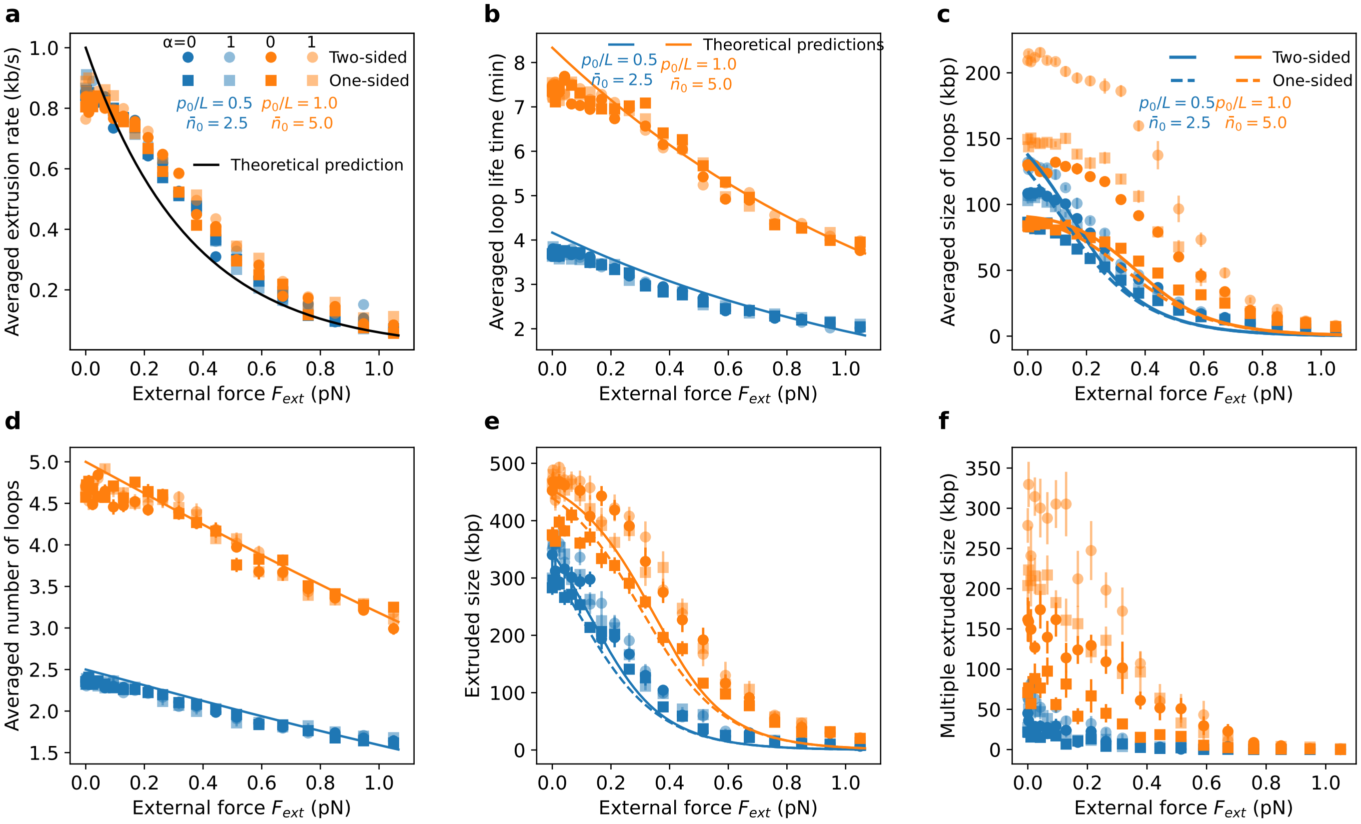


FIG. S2. Force-dependent properties of loop extrusion activity. (a) Average extrusion rate as a function of external force for various loop extrusion activities, phantom parameters, and extrusion directionalities. The black curve shows the theoretical exponential decay. (b) Average loop lifetime versus external force. Symbols are the same as in panel (a); curves represent theoretical predictions for different loop extrusion activities. (c) Average loop size as a function of external force. Symbols are the same as in panel (a); curves correspond to theoretical predictions. (d) Average number of loops as a function of external force. Symbols are the same as in panel (a); curves are the same as in panel (b). (e) Average extruded length as a function of external force. Symbols and curves are the same as in panel (c). (f) Multiple extruded length (total length extruded by more than one LEF) as a function of external force. Symbols are the same as in panel (a).

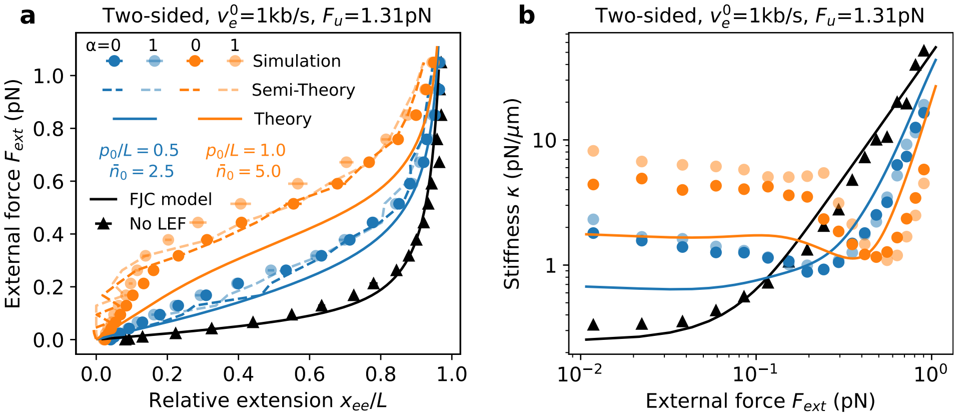


FIG. S3. Force-extension behaviour of two-sided LEFs. (a) Force-extension curves for different loop extrusion activities of two-sided LEF. Symbols represent simulation results; curves denote theoretical predictions. (b) Effective stiffness $\kappa$ as a function of external force, extracted from the force-extension data shown in panel (a). Same color legend as in (a).

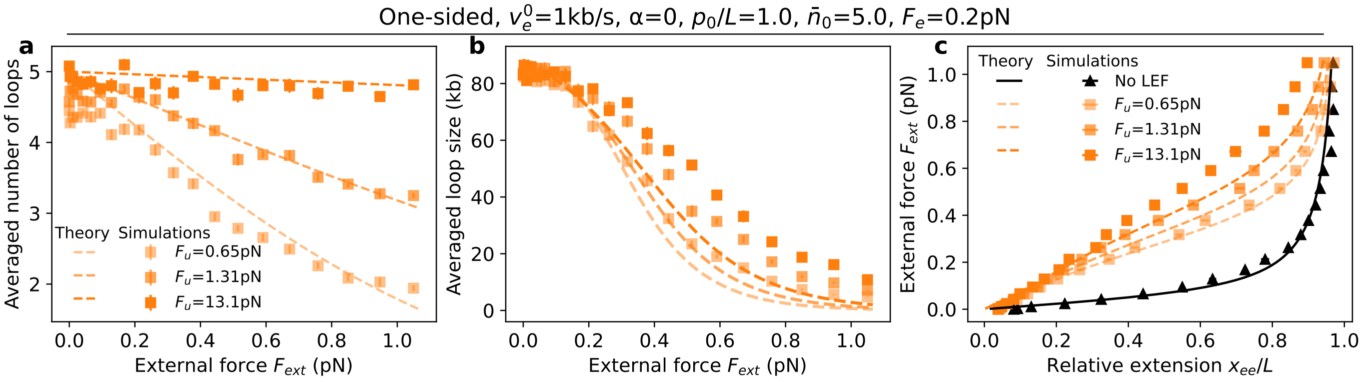


FIG. S4. Effect of the unbinding force scale $F_u$ on chromatin loop dynamics and mechanical response. (a) Average number of loops as a function of external force for different values of $F_u$. (b) Average loop size as a function of external force; the legend is the same as in panel (a). (c) Comparison of force–extension curves for varying values of $F_u$, highlighting the impact of unbinding sensitivity on chromatin mechanics.

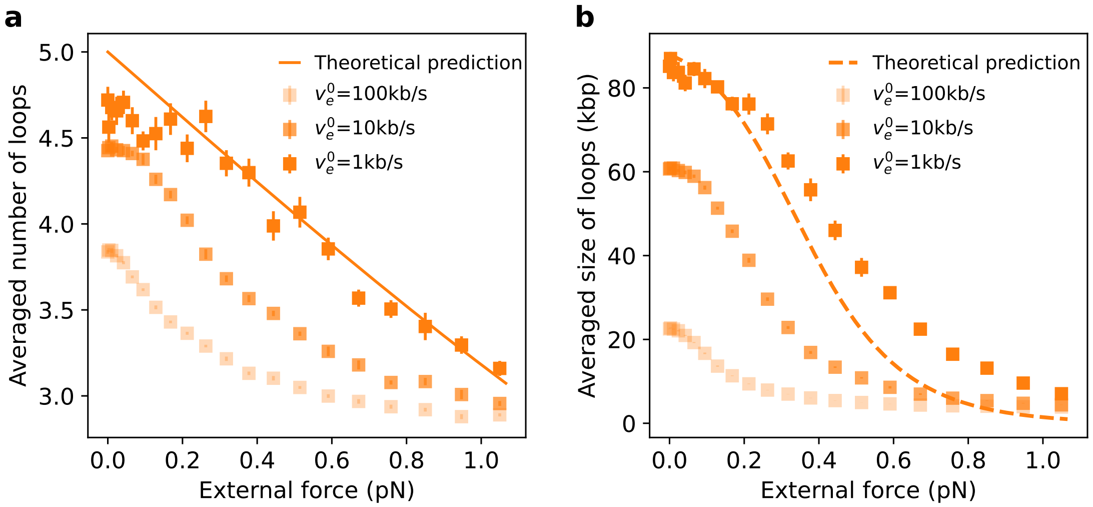


FIG. S5. Effect of extrusion speed on loop properties. (a) Average number of loops as a function of external force for one-sided loop extrusion with non-phantom extruders, $p_0/L = 1$, $\bar{n} = 5$, and different extrusion speeds. Solid curves represent theoretical predictions. (b) Average loop size as a function of external force. Symbols are the same as in panel (a), and dashed curves show the corresponding theoretical predictions. These results indicate that the loop extrusion speed significantly influences both the number and size of extruded loops under force.

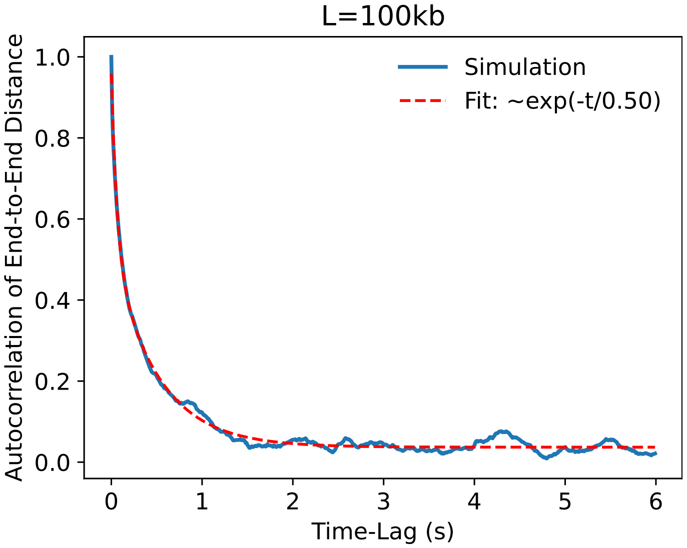


FIG. S6. Autocorrelation of the end-to-end distance as a function of time lag for a 100-kb domain embedded within a 500-kb DNA chain in the absence of loop-extrusion factors. The simulation data are fitted with a double-exponential model, yielding a longest relaxation time of approximately 0.50 s.

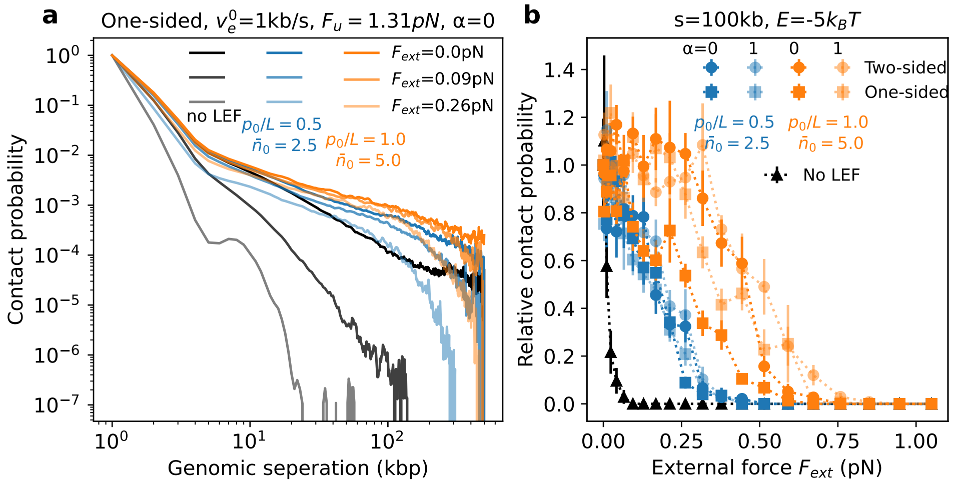


FIG. S7. Polymer structure stabilization by loop extrusion. (a) Contact probability as a function of genomic separation for different loop extrusion activities and external forces. (b) Relative contact probability between two sticky monomers separated by 100 kb, plotted as a function of external force.

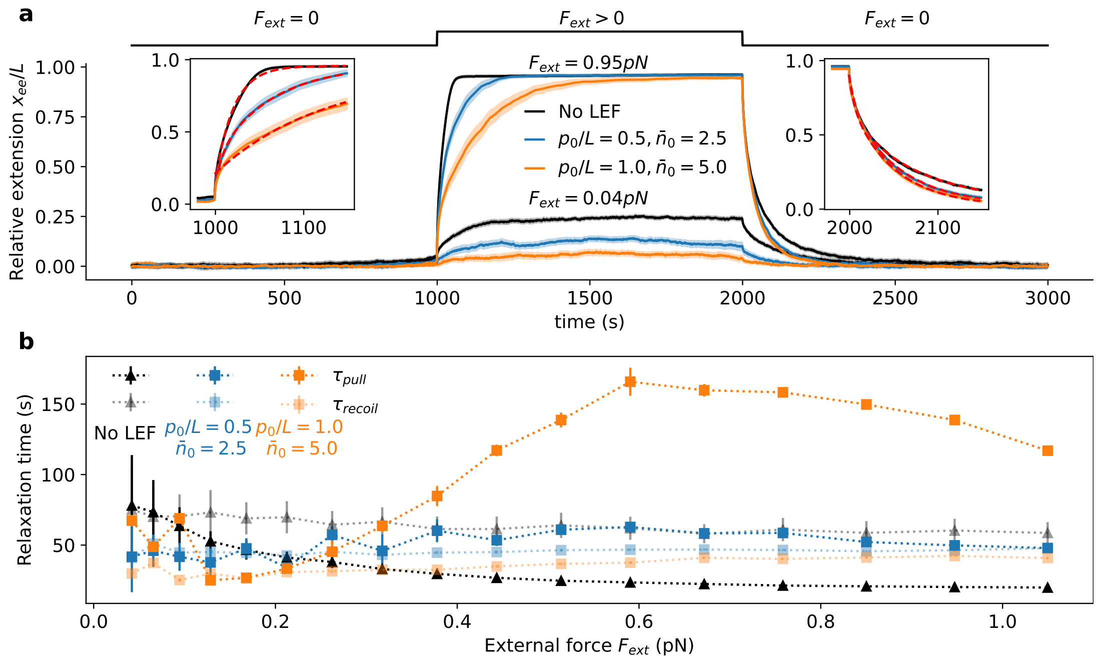


FIG. S8. Dynamic response of a polymer with loop extrusion under external force. (a) Relative extension as a function of time for different levels of loop extrusion activity. The external force is held at zero from 0 to 1000 s, then suddenly increased at $t_1 = 1000$s and maintained constant until $t_2 = 1000$s, after which it is abruptly set back to zero and held from 2000 to 3000 s. Two representative force magnitudes are shown: low force (0.04 pN) and high force (0.95 pN). Insets highlight the extension dynamics during the transition periods in the high-force case, with exponential fits shown as red dashed lines. (b) Characteristic relaxation times during pulling and release phases, obtained from exponential fits, plotted as a function of applied force for varying loop extrusion activities.

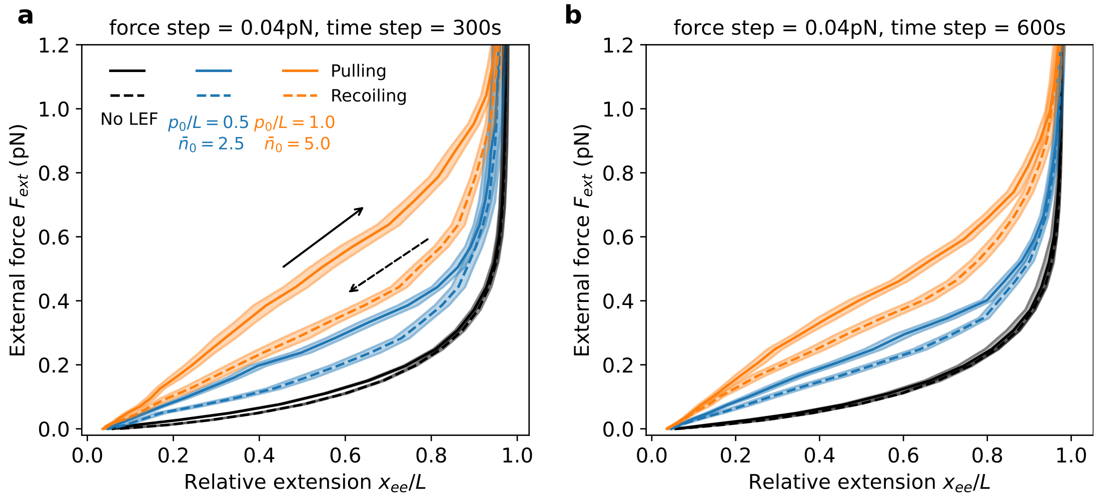


FIG. S9. Force–extension curves during pulling and recoiling under different loop-extrusion activities for (a) fast and (b) slow dynamics. The characteristic hysteresis observed in the fast case is markedly reduced for slow dynamics, consistent with its nonequilibrium origin.
